# A new approach to measure forces at junction vertices in an epithelium

**DOI:** 10.1101/2021.06.18.448930

**Authors:** Clémentine Villeneuve, Samuel Mathieu, Emilie Lagoutte, Bruno Goud, Philippe Chavrier, Jean-Baptiste Manneville, Carine Rossé

## Abstract

The mechanical properties of cell-cell junctions are critical for the stability of an epithelium. Cell-cell junction ablation experiments are classically used as a readout for junctional mechanics. However, without the knowledge of the viscoelastic properties of the microenvironment of the ablated junction, tensile junctional forces cannot be measured. Here we combine laser ablation with intracellular microrheology and develop a model to measure tensile forces exerted on cell-cell junctions. We show that the overexpression of the proto-oncogene atypical Protein Kinase C iota (aPKCi) in a single cell within a normal epithelium induces a gradient of junctional tension with neighbouring cells. Our method allows us to demonstrate that junctions contacting the aPKCi-overexpressing cell display a mechanical asymmetry which correlates with the levels of E-cadherin and P-MLC2. Measuring intracellular viscoelasticity is crucial for accurate measurements of cell-cell junction mechanics in the context of development or cancer research.

**Teaser:** We combine cytoplasmic and junctional mechanics to detect asymmetry in forces exerted on cell-cell junctions.

## Introduction

Epithelia are physiological barriers that ensure tissue integrity. The organization and maintenance of an epithelium requires adhesion complexes present along the lateral cell membranes, the so-called cell-cell junctions, and at locations where several cells contact each other, the so-called multicellular junctions. During morphogenesis or tumour progression, epithelial cells are subjected to mechanical stresses and deformations which induce junction remodelling (*1*). Cellular junctions are dynamic structures that transmit cell-generated forces through the plasma membrane to neighbouring cells (*2*). In epithelial cells, the major transmembrane protein mediating cell-cell adhesion is E-cadherin (*3*). E-cadherin adhesion promotes local actin assembly and the subsequent recruitment and activation of non-muscle myosin II at cellular junctions could enhance contractility at the plasma membrane, and thereby increase junctional tension (*3, 4*). Such mechanical forces can be sensed at cell-cell junctions by adhesion complexes, suggesting that tensile junctional forces provide a physical mode of cell–cell communication (*1, 5*–*8*). In support of this hypothesis, stresses can propagate within an epithelium on length scales of 5 to 10 cell diameters in typically one hour (*9*).

Effort has been directed towards understanding the mechanical properties and cellular behaviours that determine how an epithelium responds to strain. Multicellular junctions may adapt their mechanical properties to changes in tension and in turn respond by regulating the mechanical properties of the lateral cell-cell junctions (*10*). Quantifying the forces applied at multicellular junctions is thus critical to understand how cell-cell junctions respond to stress. Junction mechanics has primarily been studied using laser ablation experiments. In these experiments, a lateral junction between two cells is severed, which relaxes the tension exerted on the two corresponding multicellular junctions. As a consequence, the two multicellular junctions move away from each other. Tracking the locations of the two multicellular junctions over time, also known as the two vertices, allows to measure the recoil velocity, from which tensile forces are inferred (*11*). However, assuming as widely accepted that repair mechanisms such as plasma membrane repair are negligible at the time scale of laser ablation experiments, the vertices relax in the cell cytoplasm and not only experience the elastic forces exerted by the remaining junctions but also friction from the cytoplasm. Recent advances in live-cell microscopy, laser ablation and microrheology have opened the door to more quantitative approaches of tensile forces (*12*). However, to our knowledge, no method allows the direct measurement of the actual forces applied at cell-cell junctions.

We show here that microrheology experiments using intracellular optical micromanipulation combined with laser ablation allows quantification of the forces applied at multicellular junctions. We input the recoil velocity measured by laser ablation and the cytoplasm viscosity measured by microrheology into a viscoelastic model of the multicellular junction to deduce the tensile force. As a proof of concept, we took advantage of our recent study suggesting an increase in cell tension at the interface between a breast epithelial cell overexpressing the polarity protein atypical Protein Kinase C iota (aPKCi) and a surrounding wild-type (WT) epithelial cell (*13*). In this study, we showed that the effect of aPKCi overexpression depends on myosin II activity and is associated with a decrease in vinculin concentration at the cell junctions (*13*). The *PRKCI* gene coding for aPKCi is an oncogene, in particular in breast cancer, and the expression level and localization of aPKCi correlate with tumour aggressiveness (*14, 15*). Here we show and quantify how the presence of an aPKCi-overexpressing cell in a normal epithelium affects the forces applied at cell-cell junctions.

## Results

### Tension is increased at the junction between an aPKCi-overexpressing cell and surrounding wild-type cells

We have used heterogeneous MCF-10A cell monolayers to study the role of the presence of a single cell overexpressing the proto-oncogene aPKCi in regulating tensile forces within an epithelium. Cells expressing GFP-aPKCi (denoted as aPKCi cells in the following) or GFP as a control (denoted as GFP cells in the following) were mixed with wild-type (WT) cells at a low (1:100) ratio (Fig. 1A and Methods). In this study, we analyse junctional and intracellular mechanics in the context of one aPKCi cell surrounded by WT cells. Laser ablation experiments were performed on cell-cell junctions surrounding the aPKCi cell to measure the initial recoil velocity and the characteristic retraction time of the junction following ablation, two parameters classically used in ablation experiments to characterize junctional forces (*11*) (Fig. 1B, Fig. 2A and Methods). We found that junctions between an aPKCi cell and surrounding WT cells (which we call aPKCi/WT1 junctions in the following, see Fig. 1A) recoiled faster than junctions surrounding GFP cells (GFP/WT1 junctions) as indicated by an elevated initial recoil velocity and a lower retraction characteristic time (Fig. 2B). In contrast, no significant difference was observed in homogeneous monolayers of 100 % aPKCi cells or 100% GFP cells (Fig. S1A, B), These results suggest that the expression of aPKCi in a heterogeneous monolayer increases junctional tension around the aPKCi cell. We next asked whether the increase in junctional tension could propagate to WT cells further away from the aPKCi cell. We measured the initial recoil velocity and characteristic time of WT1/WT1, WT1/WT2, and WT2/WT2 junctions separating cells at increasing distances from the aPKCi cell (Fig. 1A, Fig. 2C). We found that the recoil velocity decreases while the characteristic time increases for junction further from the aPKCi cell (Fig. 2C). Both parameters reach the values measured in GFP/WT1 junctions (Fig. 2B) or in cell-cell junctions of homogeneous monolayers of 100 % aPKCi cells or 100% GFP cells (Fig. S1A, B). These results show that the increase in junctional tension induced by aPKCi expression propagates in the heterogeneous monolayer up to distances of about two cells away from the aPKCi cell.

**Figure 1.**
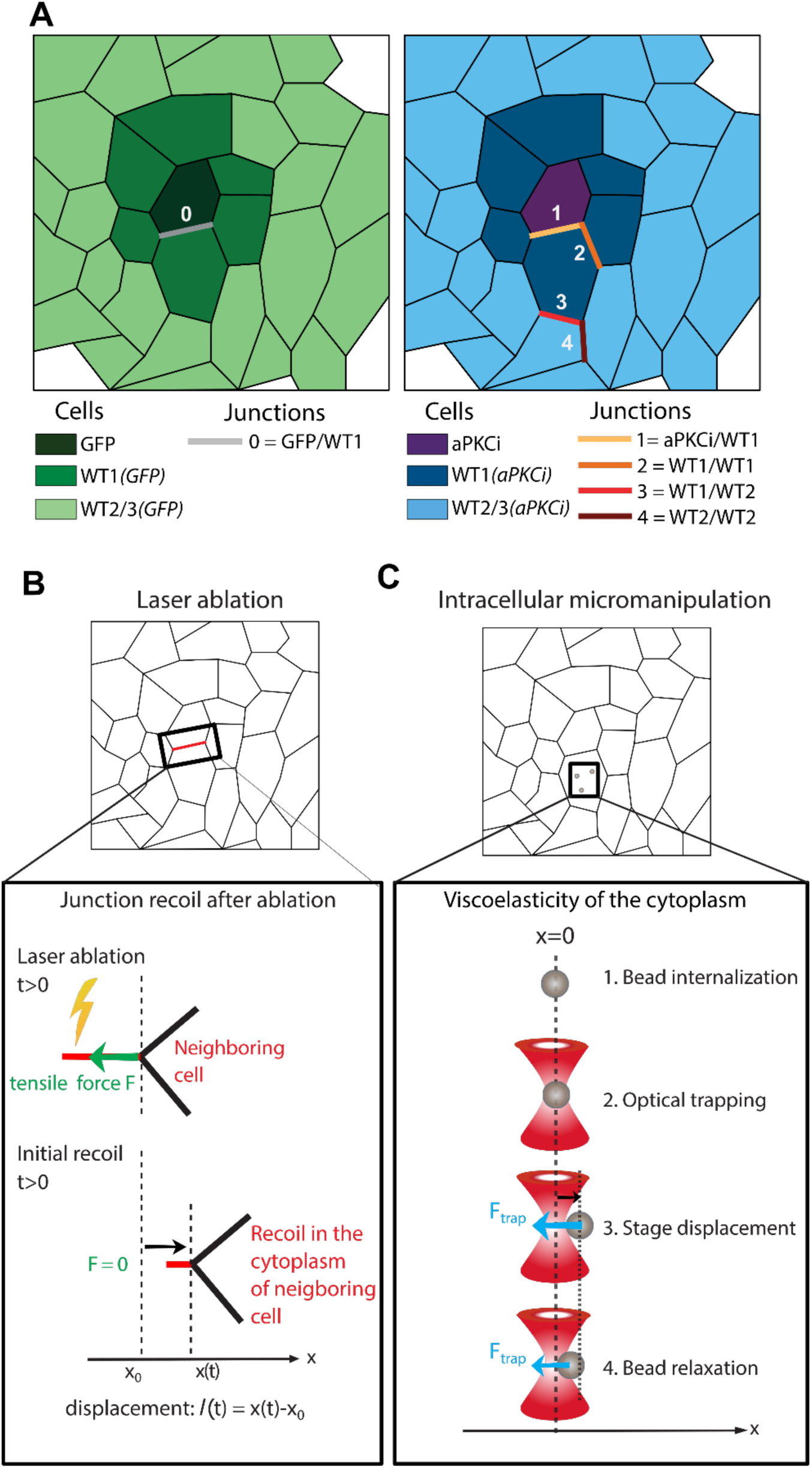
Combining laser ablation and intracellular microrheology to measure junctional tensile forces in heterogeneous cell monolayers. (**A**). Schematics defining the names given to the different cell populations and cellular junctions when control GFP cells (left) or GFP-aPKCi cells are mixed (right) with a population of wild-type (WT) cells. (**B**)-(**C**). Schematics depicting laser ablation (B) and intracellular microrheology (C) experiments. Measurements were performed on confluent cells. For heterogeneous cell populations, GFP or GFP-aPKCi were mixed with WT cells at a ratio 1:100. For laser ablation experiments (B), the vertex displacement was tracked after ablation (at *t* = 0) and the initial recoil velocity was calculated. For intracellular microrheology experiments (C), fluorescent microbeads were internalized in the cell cytoplasm and manipulated using optical tweezers. The viscoelastic relaxation of the bead towards the center of the optical trap following a step displacement of the microscope stage was measured.

**Figure 2.**
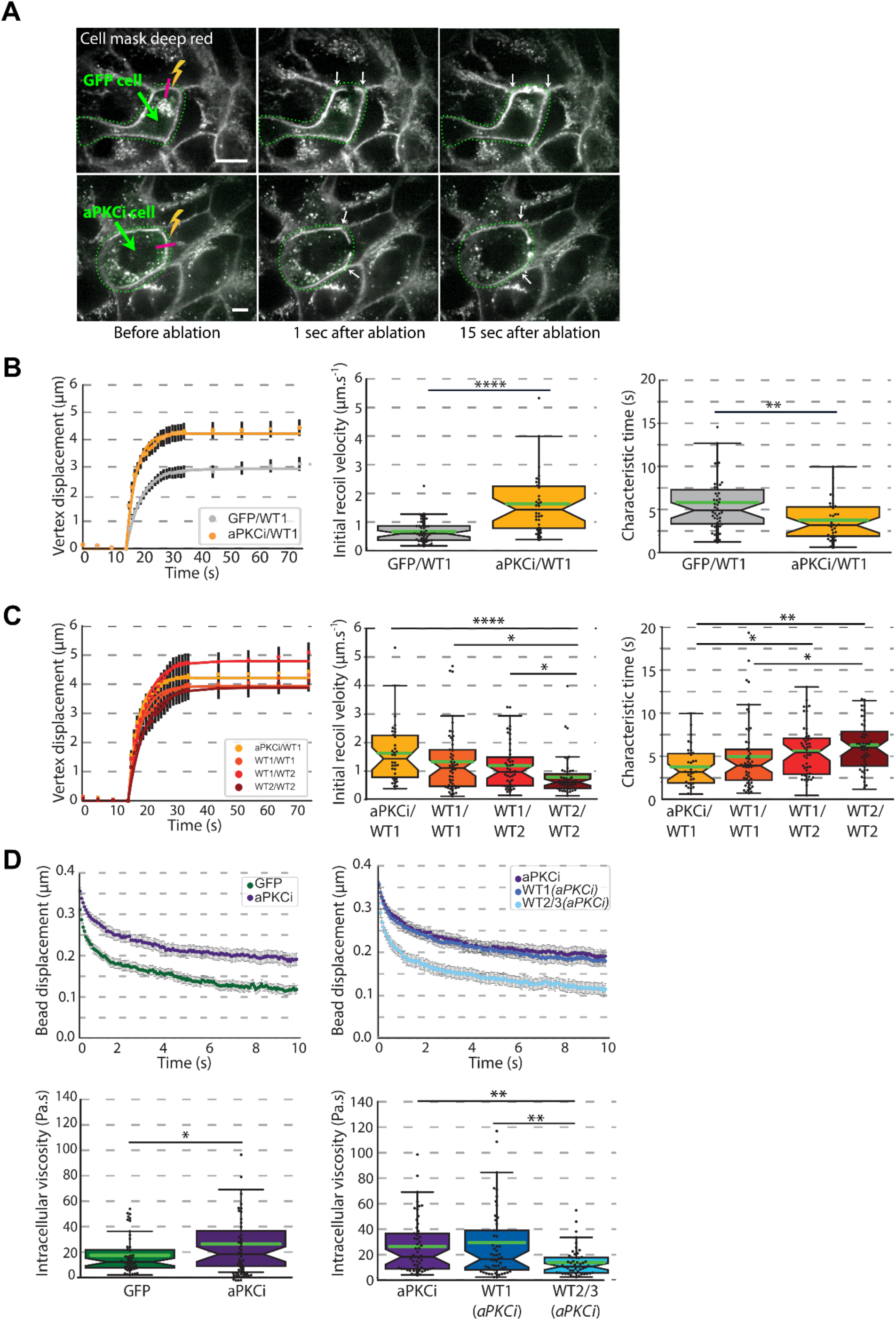
aPKCi-overexpressing cells show faster junction recoil after ablation and higher intracellular viscosity compared to surrounding wild-type cells in heterogeneous cell monolayers. (**A**). Typical laser ablation experiments in heterogeneous MCF-10A monolayers. Images of a MCF-10A GFP cell (green, top panels) and a GFP-aPKCi cell (green, bottom panels) surrounded by MCF-10A WT cells, before and after photoablation. The cell plasma membrane was labeled with Cell Mask Deep Red (grey). The ablated junction is indicated by a purple line and the corresponding relaxing vertices are shown with white arrows. The green dotted lines highlight the GFP and GFP-aPKCi expressing cells. Scale bars, 20 µm. (**B**). (Left) Vertex displacement curves after photoablation of GFP/WT1 (grey) and aPKCi/WT1 (yellow) junctions. Quantification of the initial recoil velocity (middle) and of the characteristic time (right) of the same junctions. Data are from n=46, and n=34 for GFP/WT1 and aPKCi/WT1 junctions respectively, from three independent experiments. Student’s t-tests were performed; **p< 0.01 and ****p<0.0001. (**C**). (Left) Vertex displacement curves after photoablation of aPKCi/WT1 (yellow), WT1/WT1 (orange), WT1/WT2 (red) and WT2/WT2 (dark red) junctions. Quantification of the initial recoil velocity (middle) and of the characteristic time (right) of the same junctions. Data are from n=34, 50, 48 and 48 aPKCi/WT1, WT1/WT1, WT1/WT2 and WT2/WT2 junctions respectively, from three independent experiments. Kruskal-Wallis tests were performed; *p< 0.05, **p< 0.01 and ****p<0.0001, non-significant if not specified. (**D**). Viscoelastic relaxation experiments in the cell cytoplasm. (Top) Averaged relaxation curves of the bead position following a step displacement of the stage in GFP and aPKCi cells (left), and in aPKCi cells, WT1(*aPKCi*) cells (i.e. WT cells surrounding an aPKCi cell) and WT2/3(*aPKCi*) cells (i.e. WT cells separated from an aPKCi cell by one or two WT cells) (right). (Bottom) Quantification of the intracellular viscosity of GFP and aPKCi cells (left) and of aPKCi, WT1(*aPKCi*) and WT2/3(*aPKCi*) cells (right). Data are from n=51, 53, 56 and 51 GFP, aPKCi, WT1(*aPKCi*) and WT2/3(*aPKCi*) cells respectively, from three independent experiments. A Student’s t-test (bottom left) and a Kruskal-Willis test (bottom right) were performed; *p< 0.05, and **p< 0.01, non-significant if not specified.

### Intracellular viscosity is increased in an aPKCi-overexpressing cell surrounded by wild-type cells

Because the relaxation of an ablated junction should depend on the viscosity of the cytoplasm of the cell in which the junction relaxes, we asked whether aPKCi overexpression could have an impact on intracellular viscosity of neighboring cells. We used viscoelastic relaxation experiments of internalized beads trapped in optical tweezers following a step displacement of the cell (Fig. 1C). Intracellular viscosity and elasticity were deduced by fitting the relaxation curves with the standard linear liquid (SLL) viscoelastic model (see Methods and Fig. S2A). We found that the aPKCi cells within mixed monolayers of aPKCi and WT cells exhibit significantly higher intracellular viscosity than GFP expressing cells within heterogeneous monolayers of GFP and WT cells (Fig. 2D). As in laser ablation experiments, no difference was observed in intracellular viscosity in cells in homogeneous monolayers of 100 % aPKCi cells or 100% GFP cells (Fig. S1C), further suggesting that heterogeneity in the monolayer is essential for aPKCi to induce an increase in intracellular viscosity in the neighbouring cells. Within heterogeneous monolayers of aPKCi and WT cells, WT cells in direct contact with the aPKCi cell (so called WT1(*aPKCi*) cells, Fig. 1A) exhibit the same high level of viscosity as the aPKCi cell itself (Fig. 2D). In contrast, WT cells further away (WT2/3(*aPKCi*) cells) have a lower viscosity, in the same range as that of GFP cells within mixed monolayers of GFP and WT cells (Fig. 2D) and in cells in homogeneous monolayers of aPKCi cells or GFP cells (Fig. S1C). No change in intracellular viscosity was observed in control heterogeneous monolayers of GFP and WT cells, with WT1(*GFP*) or WT2/3(*GFP*) exhibiting the same viscosity as GFP cells (Fig. S2B). The intracellular elasticity deduced from the SLL viscoelastic model followed the same trend as that of viscosity in all the conditions tested. Intracellular elasticity was higher in aPKCi cells in heterogeneous monolayers and the increased elasticity propagated to the first layer of WT cells surrounding the aPKCi cell (Fig. S2C). In contrast, control GFP expression in heterogeneous monolayers (Fig. S2B) had no effect on intracellular viscosity and elasticity. Finally, intracellular elasticity was not modified in homogeneous monolayers of GFP- or aPKCi-overexpressing cells (Fig. S1C). Together, these results show that an aPKCi cell surrounded by WT cells exhibits higher intracellular viscosity compared to control cells. Surprisingly, the increase in intracellular viscosity induced by aPKCi overexpression propagates in the heterogeneous monolayer to the WT cells in contact with the aPKCi cell.

### Combining laser ablation with microrheology shows that aPKCi overexpression increases both vertex tensile force and vertex elasticity at junctions between the aPKCi-overexpressing cell and wild-type cells

Classically, laser ablation experiments such as those presented in Figure 2B, C yield the recoil velocity *R*_0_ which reflects the junctional tension, but do not provide an actual measurement of junctional tension. During the recoil of a junction following laser ablation, the vertices of the ablated junction not only experience elastic forces from the remaining junctions but also viscous drag from the surrounding cell cytoplasm. Intracellular viscosity thus plays a role in the dynamics of junction recoil after ablation and should be taken into account to deduce junctional tensile forces from ablation experiments. Here, we have used a simple viscoelastic model for the relaxation of a vertex that includes the elastic forces exerted on the vertex which are represented by an elastic constant *K* and the friction forces applied on the vertex which are dominated by the viscosity *η* of the cytoplasm in which the ablated junction retracts (see Methods and Fig. S3A). The model allowed us to deduce the force initially applied by the junction at one vertex *F*_0_, the so-called tensile force, and the vertex elasticity *K* from the values of the initial recoil velocity *R*_0_, the characteristic time *τ* measured by laser ablation and the value of the cytoplasm viscosity *η* measured by intracellular microrheology, using *F*_0_ = *R*_0_ *g η* and 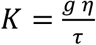, where *g* = 8*δ* if we assume that the junction relaxes as a disk of diameter *δ* (*12*) (Fig. S3B). Image analysis of relaxing junctions showed that the geometrical parameter does not vary significantly upon overexpression of aPKCi (see Methods and Fig. S4A). Consistent with the increased recoil velocity and decreased characteristic relaxation time of aPKCi/WT1 junctions (Fig. 2B) and with the increased viscosity of the aPKCi cell cytoplasm (Fig. 2D), the tensile force *F*_0_ and the vertex elasticity *K* were significantly higher for vertices of aPKCi/WT1 junctions compared to GFP/WT1 junctions (Fig. 3A). Our approach combining laser ablation and microrheology thus allows us to quantify tensional tensile forces at vertices and provides a new tool to study mechanical stress propagation in an epithelium.

**Figure 3.**
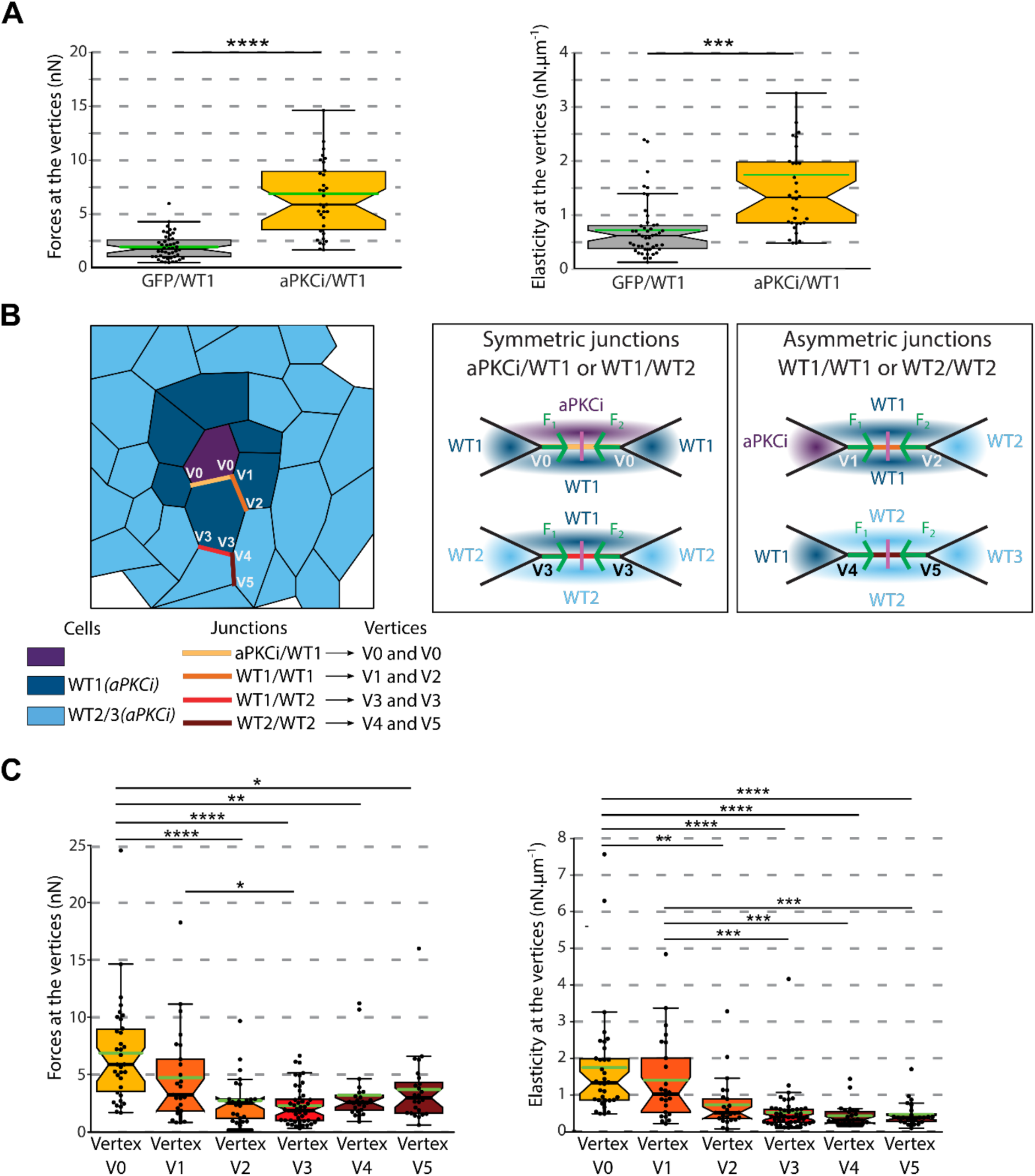
Measurements of vertex tensile force and elasticity at multicellular junctions reveals junctional asymmetry in heterogeneous cell monolayers. (**A**). Quantification of forces (left) and elasticities (right) at the vertices of GFP/WT1 and aPKCi/WT1 junctions (see Materials and Methods). Data are from n=46, and n=34 for GFP/WT1 and aPKCi/WT1 junctions respectively, from three independent experiments. Student’s t-tests were performed; ***p<0.001 and ****p<0.0001. (**B**). (Left) Schematics showing the names given to the different cells, junctions and vertices when GFP-aPKCi cells are mixed with a population of wild-type (WT) cells. (Middle) Shematics showing symmetric junctions (aPKCi/WT1 and WT1/WT2 respectively) for which the two vertices (called V0 for the aPKCi/WT1 junction and V3 for the WT1/WT2 junction) are equivalent as they both recoil in the cytoplasm of similar neighboring cells (a WT1 cell and a WT2 cell respectively). (Right) Schematics showing asymmetric junctions (WT1/WT1 and WT2/WT2 respectively) for which the two vertices (called V1 and V2 for the WT1/WT1 junction, and V4 and V5 for the WT2/WT2 junction) are not equivalent as they recoil in the cytoplasm of different neighboring cells (an aPKCi or a WT2 cell and a WT1 or a WT3 cell respectively). (**C**). Quantification of forces (left) and elasticities (right) at the vertices of symmetric or asymmetric junctions in a heterogeneous cell monolayer where aPKCi cells are mixed with WT cells. For symmetric junctions, data from the two vertices (V0 for aPKCi/WT1 junctions or V3 for WT1/WT2 junctions) were pooled. For asymmetric junctions, the two vertices were distinguished and measured separately: V1 and V2 for WT1/WT1 junctions (vertex V1 recoils towards the aPKCi cell while vertex V2 recoils towards WT2 cell); and V4 and V5 for WT2/WT2 junctions (vertex V4 recoils towards the WT1 cell while vertex V5 recoils towards the WT3 cell). Data are from n=34, 25, 25, 48, 24 and 24 V0, V1, V2, V3, V4 and V5 vertices respectively, from three independent experiments. Kruskal-Willis tests were performed; *p<0.05, **p<0.01, ***p<0.001, and **** p<0.0001, non-significant if not specified.

### aPKCi overexpression induces an asymmetry in vertex tensile force and elasticity in the proximal junction between wild-type and aPKCi-overexpressing cells

To investigate epithelial cell mechanics deeper, we applied this analysis to measure the vertex tensile force and elasticity at junctions at increasing distance from an aPKCi cell in a heterogeneous monolayer, namely the aPKCi/WT1, WT1/1WT1, WT1/WT2 and WT2/WT2 junctions (Figs. 1A and 3B). In the case of asymmetric junctions, ie. junctions for which the two vertices are not equivalent, such as WT1/WT1 and WT2/WT2 junctions (Fig. 3B), we discriminated between the two vertices since they recoil in the cytoplasm of different cells which may have different intracellular viscosity, as shown in Fig. 2D. The two vertices of WT1/WT1 junctions and of WT2/WT2 junctions have similar geometrical parameters *g* and initial recoil velocities *R*_0_ (Fig. S4A). However, for WT1/WT1 junctions, we found that the vertex tensile force and elasticity were higher for the vertex linked to the aPKCi cell (vertex V1) than for the vertex linked to the WT2 cell (vertex V2) (Figs. 3C and S4B), showing that the WT1/WT1 junction is mechanically asymmetric. In contrast, no difference was detected between the vertices (V4 and V5) of the WT2/WT2 junction (Figs. 3C and S4B), suggesting that the junctional asymmetry induced by aPKCi overexpression propagates significantly up to about two WT cells away from the aPKCi cell in the heterogeneous monolayer.

### The levels of P-MLC2 and E-cadherin along asymmetric junctions correlate with the vertex tensile force and elasticity

We have showed previously that aPKCi overexpression induces a decrease in vinculin at cell-cell junctions associated with an increase junctional tension when the aPKCi cell is surrounded by WT cells (*13*). To test whether the asymmetry in vertex tensile force and elasticity that we evidenced in junctions around the aPKCi cell in a heterogeneous monolayer reflects an asymmetry in cell-cell adhesion and contractility, we have investigated the distribution of E-cadherin and P-MLC2 along WT1/WT1 junctions (Fig. 4). We found that the levels of P-MLC2 and E-cadherin are significantly increased at the junction between the aPKCi cell and the two WT1 cells (i.e. at vertex V1) compared to the junction between the WT2 cell and the two WT1 cells (i.e. at vertex V2) (Fig. 4A and B). These results show that the higher levels of E-cadherin and P-MLC2 correlate with elevated vertex tensile force and elasticity. We propose that the gradient of vertex tensile force and elasticity around the aPKCi cell originates from a gradient in cell-cell adhesion and junction contractility (Fig. 4C).

**Figure 4.**
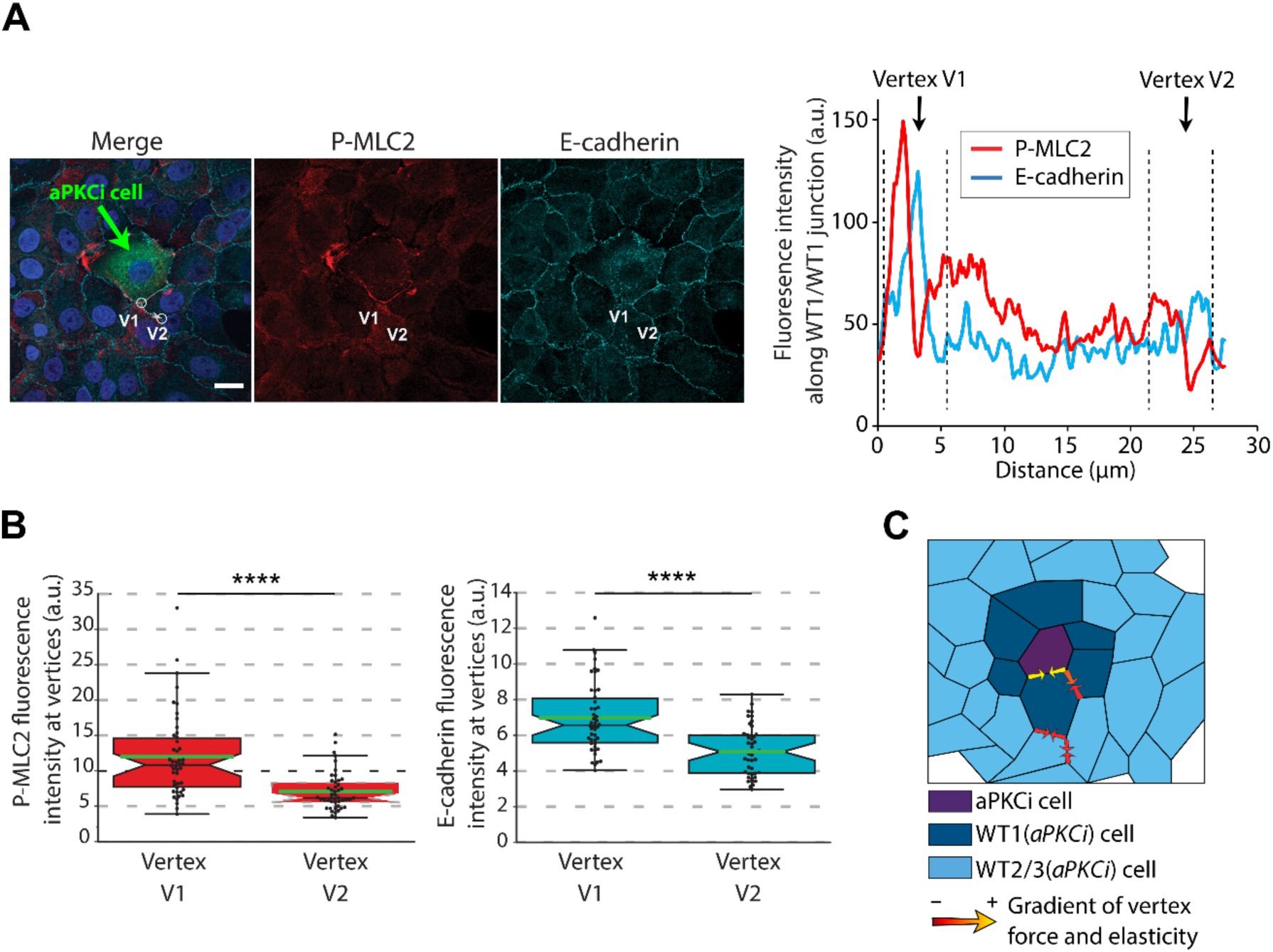
Asymmetric tensile forces along heterogeneous cell-cell junctions are associated with asymmetric levels of P-MLC2 and E-cadherin. (**A**). (Left) Representative confocal images of a MCF-10A aPKCi cell surrounded by MCF-10 WT cells stained for P-MLC2 (red), E-cadherin (cyan) and nuclei (DAPI, blue). Scale bar, 20 µm. (Right) Fluorescence intensity profile of P-MLC2 (red line) and E-cadherin (blue line) averaged along the asymmetric WT1/WT1 junction shown in the top left panel (grey dotted line, 7 µm thick). The aPKCi cell and vertex V1 are on the left (distance ∼2.5 µm), while the WT2 cell and vertex V2 are on the right (distance ∼25 µm). (**B**). Quantification of P-MLC2 (left) and E-cadherin (right) fluorescence intensities at V1 and V2 vertices. Data are from n=46 vertices, from three independent experiments. The area quantified for each vertex is shown as white circles (7 µm diameter) in A (bottom left panel). Student t-tests were performed; ****p<0.0001. (**C**).Schematics showing how vertex forces and elasticity are distributed around an aPKCi cell in a heterogeneous monolayer. Combining laser ablation with intracellular microrheology allows to detect an increase in vertex forces and elasticities towards the aPKCi cells, positively correlated with P-MLC and E-cadherin levels at the vertices.

## Discussion

### Measurements of tensile forces at junctions in an epithelium

Characterizing the forces applied at cell-cell junctions is crucial to understand the impact of mechanical stresses on the integrity of tissues and cell behaviour within an epithelium (*8*). Here, by combining laser ablation with intracellular microrheology and a viscoelastic model, we measure the tensile force applied at the junction between epithelial MCF10A cells in a confluent monolayer. We report values of tensile forces in the nanonewton range (2 nN up to 7nN depending on the experimental conditions). These tensile forces are generated by the cells themselves through the acto-myosin cytoskeleton (Figure 4). Because E-cadherin-based junctions couple the contractile acto-myosin cytoskeleton to the cell membrane, E-cadherin is thought to participate in the generation of junctional tension (*8*). Moreover, E-cadherin adhesion promotes the recruitment and activation of non-muscle myosin II to cell junctions (*16*). As a consequence, the junctional tensile forces that we measure in our experiments mainly result from the coupling of E-cadherin junctions to the acto-myosin cytoskeleton. Consistently, the asymmetry in the levels of E-cadherin and P-MLC2 we detect along proximal junctions between wild-type and aPKCi-overexpressing cells correlates with an asymmetry in the tensile force experienced by the corresponding vertices. A similar mechanical heterogeneity along cell-cell junctions was recently reported in the context of convergent extension in *Xenopus* embryos and was shown to be due to a patterned clustering of the Cdh3 cadherin (*17*).

### The rheological properties of aPKCi-overexpressing cells depend on the surrounding environment

One surprising finding of our study is that aPKCi overexpression only triggers changes in intracellular viscoelasticity in an aPKCi cell surrounded by WT cells. We show that both elasticity and viscosity increase in the cytoplasm of aPKCi cells seeded at very low density in a WT monolayer (Fig. 2D and Fig. S2C), while no effect is detected when aPKCi cells are cultured as a homogenous epithelium (Fig. S1C). Strikingly, elasticity and viscosity are also increased in the first WT neighbours of the aPKCi cell (WT1(aPKCi) cells in Fig. 2D and Fig. S2C). The effect is short ranged since both the elasticity and viscosity dropped to control levels in the second and third neighbours of the aPKCi cell (WT2/3(aPKCI) in Fig. 2D and Fig. S2C). A feedback between cell contractility and intracellular mechanics could explain why the rheological properties of aPKCi-overexpressing cells depend on the presence of WT cells. We have shown previously that aPKCi cells surrounded by WT cells exhibit increased contractility (*13*). Consistently, we show here that P-MLC2 levels are elevated in aPKCi cells but not in the adjacent WT cells (Fig. 4). Such an asymmetry could provide a mechanical cue to the aPKCi cell, which would induce an increase in intracellular viscosity. In agreement with this hypothesis, it was recently shown that MCF10A cells overexpressing Ras^V12^ cultured on a stiff substrate exhibit intracellular stiffening (*18*). Oncoproteins such as Ras or aPKCi, despite having different oncogenic properties, may thus modulate cell rigidity depending on the microenvironment, for instance the nature of the extracellular matrix or the mechanical properties of the surrounding cells (*13, 14, 19, 20*). Our results suggest that polarity proteins such as aPKC, which regulate cell–cell adhesion and cell contractility and are often deregulated in cancers, play crucial roles in tissue mechanics and architecture during tumorigenesis, in particular at the onset of tumour development. The presence of a transformed cell within a normal epithelium may affect local intracellular mechanics and tensile forces at the vertices of the proximal surrounding cells.

### Combining laser ablation with microrheology to detect asymmetry in vertex tensile forces

Within an epithelium, stresses have been shown to be transmitted typically over five to ten cell diameters scale on time scales of an hour (*9*). However, it is not clear if and how the presence of a modified cell in the epithelium perturbs the mechanics of the epithelium locally and can be detected at a distance from the modified cell through stress propagation. Here, by combining laser ablation and microrheology, we were able to measure the tensile forces applied at the vertices of cells surrounding a modified (overexpressing aPKCi) cell within a WT epithelium. We detect an asymmetry in vertex tensile force and elasticity of the vertices in the first raw of WT cells surrounding the aPKCi cell (Fig. 3C) which generates a gradient of tensile forces decreasing from the aPKCi cell (Fig. 4C).

In our study, we have used MCF10A monolayers as model epithelia. The geometry of such monolayers is not as regular as in other invertebrate or vertebrate model systems in which vertices are tri-cellular junctions (TCJs). Despite the complex geometry, our method provides enough resolution to detect subtle differences in tensile forces. We can thus speculate that our approach will prove useful in more organized epithelia, in particular at TCJs. Numerous functions have been attributed to TCJs in epithelial tissues, for instance in the regulation of cell division, collective cell migration and cellular mechanical properties (*21*–*23*). In normal tissues, TCJs are considered as hubs where biochemical and mechanical signalling occur to control local cell dynamics while maintaining tissue integrity (*8*).

In pathological conditions, TCJs between cancer and normal cells could also play crucial roles during tumour progression (*13, 22, 24*–*26*), especially in cases where oncogenes are not sufficient to drive tumour initiation. One speculation is that tumour progression results from an aberrant communication between cancer cells and their microenvironment (*27*)(*22, 24, 25, 27*). Alteration of tensile forces at TCJs could participate in such aberrant communication.

We anticipate that our method combining laser ablation and microrheology will allow a more thorough and precise investigation of tensile forces at vertices in several epithelial models. Optical tweezers-based microrheology can be applied not only in 2D cell monolayers, but also in 3D organoids (*28*). Although technically challenging, combining laser ablation and microrheology in model organisms could also benefit studies on development and tumour progression.

## Materials and methods

### Cell culture and stable cell lines

MCF10A cells were maintained in DMEM-F12 (#11039-021, Gibco) containing 10% Penicillin-Glutamine, 10µg/mL human insulin (#I9278, Sigma Aldrich), 100 ng/mL cholera toxin (Sigma-Aldrich), 0.5mg/mL hydrocortisone (Sigma Aldrich), 5% horse serum and EGF 20 ng/ml (PeproTeck) at in 5% CO2 incubator at 37°C. MCF10A stably expressing fluorescent reporter constructs were generated by lentiviral transduction followed by FACS sorting (*13*). The absence of mycoplasma contamination in cell cultures was routinely verified using a PCR test. When needed, doxycycline (Sigma-Aldrich) was used at a concentration of 1 µg/mL.

### 2D immunostaining of MCF10A cells

To measure E-cadherin (clone ECCD2, Invitrogen, ref. 13-1900) and pMLC2 (phospho-Serine 19, Ozyme, ref. 3671S) fluorescence intensity, MCF10A cells overexpressing doxycycline inducible GFP or GFP-aPKCi were mixed with MCF10A-WT cells at a ratio of 1:20 on glass coverslips in MCF10A medium (*13*). Doxycycline was added in the medium 12 hours later or GFP or GFP-aPKCi cells were induced 24h before seeding. Cells were first fixed with PBS-4%-PFA for 10 minutes, followed by permeabilization with PBS-0.5% Triton for 10 minutes. Blocking was performed in PBS-10% FBS for 1 hour. Primary antibody (E-cadherin diluted in 1/100; pMLC2 diluted in 1/50) incubation was performed in PBS-10% FBS for 3 hours at room temperature. Secondary antibody incubation was performed for 45 minutes in PBS. Samples were mounted in Prolong-DAPI.

### Laser ablation

MCF10A-WT and GFP or GFP-aPKCi cells were mixed at a ratio 1:100 and plated on a glass coverslip 16 hours before acquisition. GFP or GFP-aPKCi expression was previously induced with doxycycline, 24 hours before seeding. The laser ablation system was composed of a pulsed 355-nm-20kHz ultraviolet laser (Gataca System) interfaced with an iLas system running in parallel with Metamorph 7 Software. Laser ablation was performed at 37°C and 5% CO2 on a confocal spinning disk (Yokagawa CSU-X1 spinning head on a Nikon Eclipse T*i-E* inverted microscope) equipped with an EMCCD camera (Evolve, Photometrics) and a 100x oil immersion objective (Nikon S Fluor 100x 0.5-1.3 NA). The ablation region was drawn as a line crossing perpendicularly the middle of the junction at the interface between a MCF10A-WT cell and a GFP or GFP-aPKCi overexpressing MCF10A cell. Images were acquired every 5 seconds during 15 seconds before ablation. For photoablation, the laser beam was focused on the region of interest during a pulse of 80-90 ms at 85% laser power. After photoablation, acquisitions were performed every 1 second for 20 seconds and then every 10 seconds for another 2 minutes to capture the full displacement of the two vertices. The displacement between the two vertices *L*(*t*) relative to their initial positions, was then tracked using ImageJ and the speed of retraction *v*_*ablation*_ (termed “initial recoil velocity”) and the characteristic time *τ*_*ablation*_ were obtained after fitting the displacement curve *L*(*t*) with a one-phase association exponential using GraphPad Prism (GraphPad Software).

### Intracellular optical micromanipulation

The set-up combining optical trapping and fast confocal imaging has been described previously (*29, 30*). In brief, doxycycline inducible MCF10A cells overexpressing GFP or GFP-aPKCi were mixed with MCF10A-WT cells at a ratio of 1:100 on glass-bottom culture dishes (MatTek P50G-1.5-14-F) in MCF10A medium. GFP or GFP-aPKCi cells were induced 24h before seeding. Red fluorescent (exc. 580/em. 605nm) 2μm diameter latex beads (ThermoFisher F88265) were endocytosed overnight. We have showed previously that the beads are enclosed in a lysosomal membrane closely apposed on the bead surface which does not perturb the mechanical measurements (*29*). The incubation time and bead concentration were adjusted so that cells typically contained one or two beads before optical micromanipulation. Immediately prior to the experiment, the culture medium was supplemented with 20 mM Hepes. A bead located in the cytoplasm in the perinuclear region was first trapped using the optical tweezers and a small (∼0.2 *μm*) step displacement of the stage was performed to detach the bead from potential links to the cytoskeleton. The microscope stage was then displaced in order to push the surrounding cytoplasm against the bead. The applied friction force on the bead *F*_*trap*_ was deduced from the bead displacement relative to the center of the trap Δ*x*, after calibration of the trap stiffness using Stokes law in water *k*_*trap*_ = 234 ± 40 *pN. μm*^−1^:

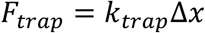

Stage displacement was performed using a nanopositioning piezo-stage (Nanobio 200, Mad City Labs) controlled by the NanoRoute3D software (Mad City Labs). The displacement consisted of one 0.5 *μm* step followed by a 10 *s* pause to allow visco-elastic relaxation of the bead towards the trap center. The total duration of optical trapping was limited to 30 *s* for a given cell to ensure cell viability (output power of the infra-red trapping laser at the objective aperture was 150 *mW*). The position of the center of the bead relative to the center of the trap was tracked using a homemade Matlab code.

### Viscoelastic model of the cytoplasm

The movement of the bead (radius *R* = 1*μm*) in the cytoplasm is governed by the coupling between the mechanical properties of the cell interior and the optical trap. The bead is subjected to the visco-elastic forces due to the deformation of the microenvironment around it and to the force exerted by the optical tweezers, that act as a spring of stiffness *k*_*trap*_. The microenvironment of the bead can be modelled by a three-element visco-elastic model, known as the Standard Linear Liquid (SLL) model, composed of a Kelvin-Voigt body, which consists of a spring of stiffness *k* and a dashpot of viscosity *η* in parallel, and a dashpot of viscosity *η*_0_ in series (Fig. S2A). The position of the center of the trap is fixed at *x* = 0. The initial positions of the stage, the bead and the node between the Kelvin-Voigt body and the dashpot in series are respectively *X*_*S*_ = 0.5 *μm, X*_*b*_ and *X*_1_ (where *X*_*b*_ < *X*_*S*_ and *X*_1_ = *X*_*S*_). *X*_1_ − *X*_*b*_ is the free length of the spring of stiffness *k*. The initial position leads to a force *F*_*trap*_ exerted by the optical trap given by (projected on the *x* axis): *F*_*trap*_ = −*k*_*trap*_*x*_*b*_, where *x*_*b*_(*t*) is the position of the bead as a function of time. This force brings the bead back to the center of the trap and thus deforms the visco-elastic components modeling the cell cytoplasm. These components then provide other forces which are opposed to the movement of the bead. The forces due to the Kelvin-Voigt body can be decomposed into a sum of a viscous drag *F*_*v*_ and an elastic contribution *F*_*e*_ given by (projected on the *x* axis): 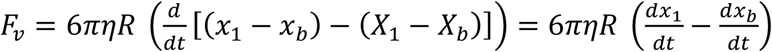 and *F*_*e*_ = *k* [(*x*_1_ − *x*_*b*_) − (*X*_1_ − *X*_*b*_)], where *x*_1_(*t*) is the position of the node between the Kelvin-Voigt body and the dashpot in series. In Newton’s law, inertia can be neglected thus leading to:

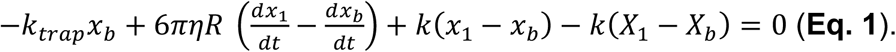

Stress continuity between the Kelvin-Voigt body and the dashpot in series imposes:

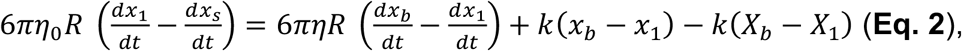

where *x*_*S*_ is the position of the stage relative to the trap center. Since the microscope stage is only submitted to a step displacement, *x*_*S*_(*t*) is a constant given by the amplitude of the step: *x*_*S*_(*t*) = *X*_*S*_ = 0.5 *μm*. Introducing this value in **Eq. 2** and combining it with **Eq. 1** yields the following equation:

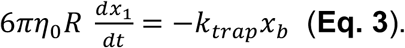

Combining **Eq. 1** and **Eq. 3** leads to the differential equation verified by

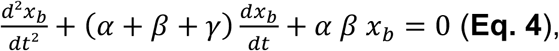

with 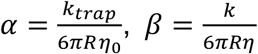 and 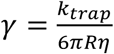. Solving this equation with the initial conditions *x*_*b*_(*t* = 0) = *X*_*b*_ and 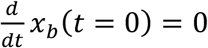 gives:

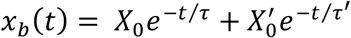

where the two relaxation times *τ* and *τ*′ satisfy (*τ*^−1^)(*τ*′^−1^) = *αβ* and *τ*^−1^ + *τ*′^−1^ = *α* + *β* + *γ* and with 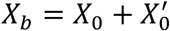. Relaxation curves giving the bead displacement relative to the trap center *x*_*b*_(*t*) can then be fitted for 0 ≤ *t* ≤ 10 *s* to yield the different parameters and finally deduce the rheological parameters *k, η* and *η*_0_ (Fig. S1B). For simplicity, we term the spring stiffness *k* (in N.m^-1^) the elasticity of the cell cytoplasm. Note that the dashpot in series of viscosity *η*_0_ dominates the response at long timescales for which the experimental data is more variable. Consequently, the value of *η*_0_ is less relevant than that of the viscosity of the Kelvin-Voigt body *η* and is not shown in the results. Nevertheless, the dashpot in series allows a better fitting of the relaxation curves and a more accurate determination of the rheological parameters of the Kelvin-Voigt body.

### Viscoelastic model to derive the vertex junctional tensile force and the vertex elasticity from laser ablation combined with intracellular microrheology

The viscoelastic model used to derive the junctional tensile force applied on one vertex (point where three junctions meet) and the vertex elasticity is depicted in Fig. S3. In a laser ablation experiment, which occurs at time *t* = 0, we assume that the forces acting on the retracting vertex are only due to the stiffness *K* of the two remaining junctions (which we term vertex elasticity for simplicity, in N.m^-1^) and to the viscous drag due to the viscosity *η* of the cytoplasm in which the ablated junction retracts (Fig. S3A). The force balance on the retracting vertex is given by 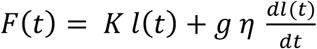, where *F*(*t*) is the force exerted by the junction to be ablated, *l*(*t*) is the position of the vertex and *g* is a geometrical parameter characterizing the relaxing junction. Assuming that the junction relaxes as a disk of diameter *δ* (Fig. S3B), the value of the geometrical parameter *g* is *g* = 8*δ* (*12*). The diameter *δ* was measured at a fixed distance *d* = 5 μm from the relaxing vertex (Fig. S3B) and was found to be similar for the two vertices of WT1/WT1 and WT2/WT2 asymmetric junctions (data not shown), yielding ratios between the geometrical parameters of the two vertices close to one (Fig. S4A). At time *t* ≤ 0, the vertex is immobile, its position is *l*_0_ and the force exerted on the vertex is the junctional tensile force *F*_0_ so that *F*_0_ = *Kl*_0_. After ablation, at time *t* > 0, the force *F*(*t*) vanishes and *l*(*t*) obeys the differential equation 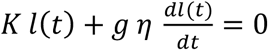 which yields 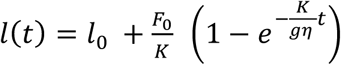.

Laser ablation experiments classically allows to determine the recoil velocity (*11*). If the vertices recoil symmetrically, the displacement between the two vertices *L*(*t*) relative to their initial positions is given by 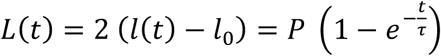, where 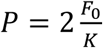 is the plateau and 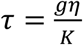 is the retraction characteristic time. The initial recoil velocity, i.e. the initial retraction speed of the vertices, is then 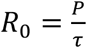.

Combining the measurements of the initial recoil velocity *R*_0_ and the characteristic time *τ* obtained from laser ablation experiments and the measurements of cytoplasm viscosity *η* obtained from intracellular microrheology experiments allows us to deduce the force initially applied by the junction at one vertex *F*_0_ (i.e. the tensile force) and the combined elasticities of the two other junctions connected with the same vertex *K* (i.e. the vertex elasticity): *F*_0_ = *R*_0_ *g η*/2 and 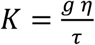. From the expressions of the tensile force *F*_0_ and the vertex elasticity *K*, an asymmetry between the two vertices of a given junction can arise from differences in the geometrical parameter *g* characterizing the geometry of the vertex retraction and/or from differences in the viscosity *η* of the medium in which the vertex is retracting. For symmetric junctions (GFP/WT1, aPKCi/WT1 and WT1/WT2, see Fig. 1A) for which *g* and *η* are the same since the two vertices recoil in the cytoplasm of similar neighboring cells, data for the two vertices were pooled. In contrast, for asymmetric junctions (WT1/WT1 and WT2/WT2, see Fig. 1A), the two vertices are not equivalent as they recoil in the cytoplasm of different neighboring cells and *g* and *η* may be different for each vertex. The two vertices were distinguished depending on the cell in which the vertex recoils (for WT1/WT1 junctions: WT1/WT1(*aPKCi*) or WT1/WT1(*WT2*) for the vertex recoiling towards the aPKCi cell or towards the WT2 cell, respectively; for WT2/WT2 junctions: WT2/WT2(*WT1*) or WT1/WT1(*WT3*) for the vertex recoiling towards the WT1 cell or towards the WT3 cell, respectively). Asymmetry was quantified by calculating the ratios between the tensile forces at the two vertices (*F*_0 WT1/WT1(*aPKCi*)_/*F*_0 WT1/WT1(*WT*2)_ for WT1/WT1 junctions and *F*_0 WT2/WT2(*WT*1)_/*F*_0 WT2/WT2(*WT*3)_ for WT2/WT2 junctions) and the ratios between the elasticities at the two vertices (*K*_WT1/WT1(*aPKCi*)_/*K*_WT1/WT1(*WT*2)_ for WT1/WT1 junctions and *K*_WT2/WT2(WT3)_/*K*_WT2/WT2(*WT*3)_ for WT2/WT2 junctions).

### Statistical analysis

Results are representative of at least 3 independent experiments. Data are expressed as means ± standard error mean. When two conditions are compared, statistical relevance was evaluated using Student’s t-tests. When more than two conditions are compared, statistical relevance was evaluated a global Kruskal-Wallis test followed by post hoc operations (Dunn’s test). p-values are indicated as non-significant (n.s.) p>0.5, * p<0.05, ** p<0.01, *** p<0.001, **** p<0.0001.

## Acknowledgements

The authors acknowledge the Nikon Imaging Center at the Institut Curie-CNRS, the PICT-IBiSA (member of the France-BioImaging national research infrastructure) for the fluorescence microscopy equipment and help with image acquisition. We are particularly grateful to the Flow Cytometry Core Facility of the Institut Curie. Members of P. Chavrier and B. Goud laboratories are thanked for helpful discussions.

## Funding

Funding for this work was provided by the following grants.

*Fondation ARC pour la Recherche contre le Cancer*, grants PJA20141201972 and PJA20181208073 (CR)

*Cancéropôle Ile de France*, grants EMERG-1 and INVADOID 2016 (CR)

GEFLUC Île-de-France - Cancer Research 2019 (PC)

LABEX Cell(n)Scale ANR-11-LABX-0038, ANR-10-IDEX-0001-02 and core funding from the *Institut Curie* and *Centre National pour la Recherche Scientifique* (CR, JBM, PC, BG). INSERM Plan Cancer 2014-2019 INSERM, grant number 17CP089_00 (JBM)

PhD fellowships from PSL Research University and *Fondation pour la Recherche Médicale* (CV)

PhD fellowship from Sorbonne Université UPMC University Paris 06, grant *Programme Doctoral “Interfaces Pour le Vivant”* (SM)

## Author contributions

Conceptualization: CR, JBM, PC, BG

Methodology: CV, SM, EL, CR, JBM

Investigation: CV, SM, EL

Supervision: CR, JBM

Writing—original draft: CR, JBM

Writing—review & editing: JBM, CR, CV, SM, PC, BG

## Data and materials availability

All data are available in the main text or the supplementary materials

## Supplementary Materials

Supplementary materials contain four supplementary figures. See corresponding file

