## Supplemental Data PDF for "A new approach to measure forces at junction vertices in an epithelium"

**This PDF file includes:**

Figs. S1 to S4

### Supplementary Figure 1

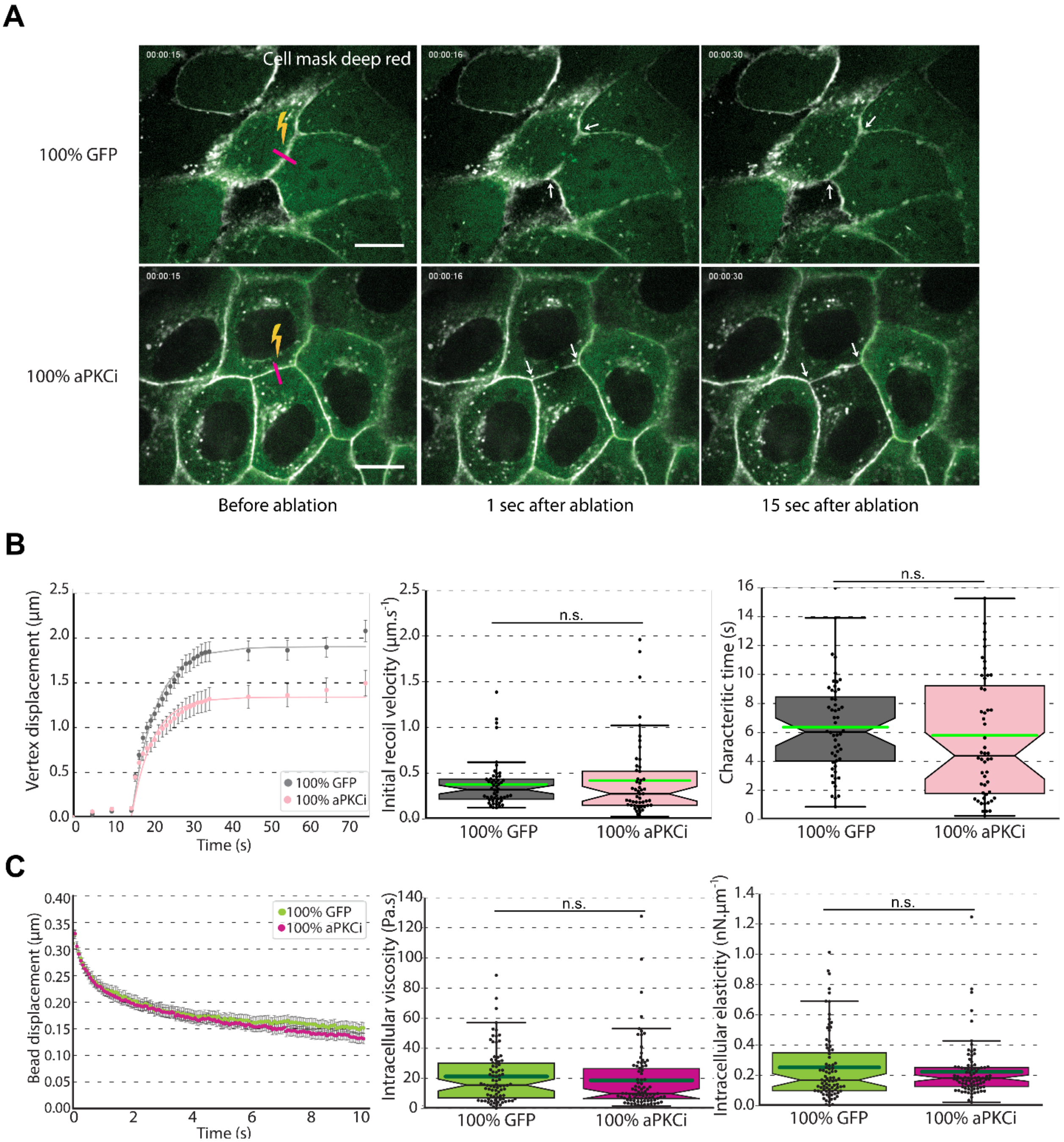

Fig. S1.
(related to Fig. 2)

**Laser ablation and intracellular microrheology experiments in homogeneous cell monolayers of MCF-10A cells expressing either GFP (100% GFP) or aPKCi (100% aPKCi).**

(**A**). Typical laser ablation experiments in homogeneous MCF-10A monolayers. Images of MCF-10A 100% GFP (top panels) and 100% aPKCi (bottom panels) cell monolayers, before and after photoablation. The cell plasma membrane was labeled with Cell Mask Deep Red (grey). The ablated junction is indicated by a purple line and the corresponding relaxing vertices are shown with white arrows. Scale bars, 20 µm.

(**B**). (Left) Vertex displacement curves after photoablation. Quantification of the initial recoil velocity (middle) and of the characteristic time (right). Data are from n=52 and n=52 junctions from 100% GFP and 100% aPKCi monolayers respectively, from three independent experiments. Student's t-tests were performed; ns, non-significant.

(**C**). (Left) Averaged viscoelastic relaxation curves of the bead position following a step displacement of the stage in cells from 100% GFP and 100% aPKCi monolayers (left). Quantification of the intracellular elasticity (middle) and viscosity (right). Data are from n=84 and n=86 cells from 100% GFP and 100% aPKCi monolayers respectively, from three independent experiments. Student's t-tests were performed; ns, non-significant.

### Supplementary Figure 2

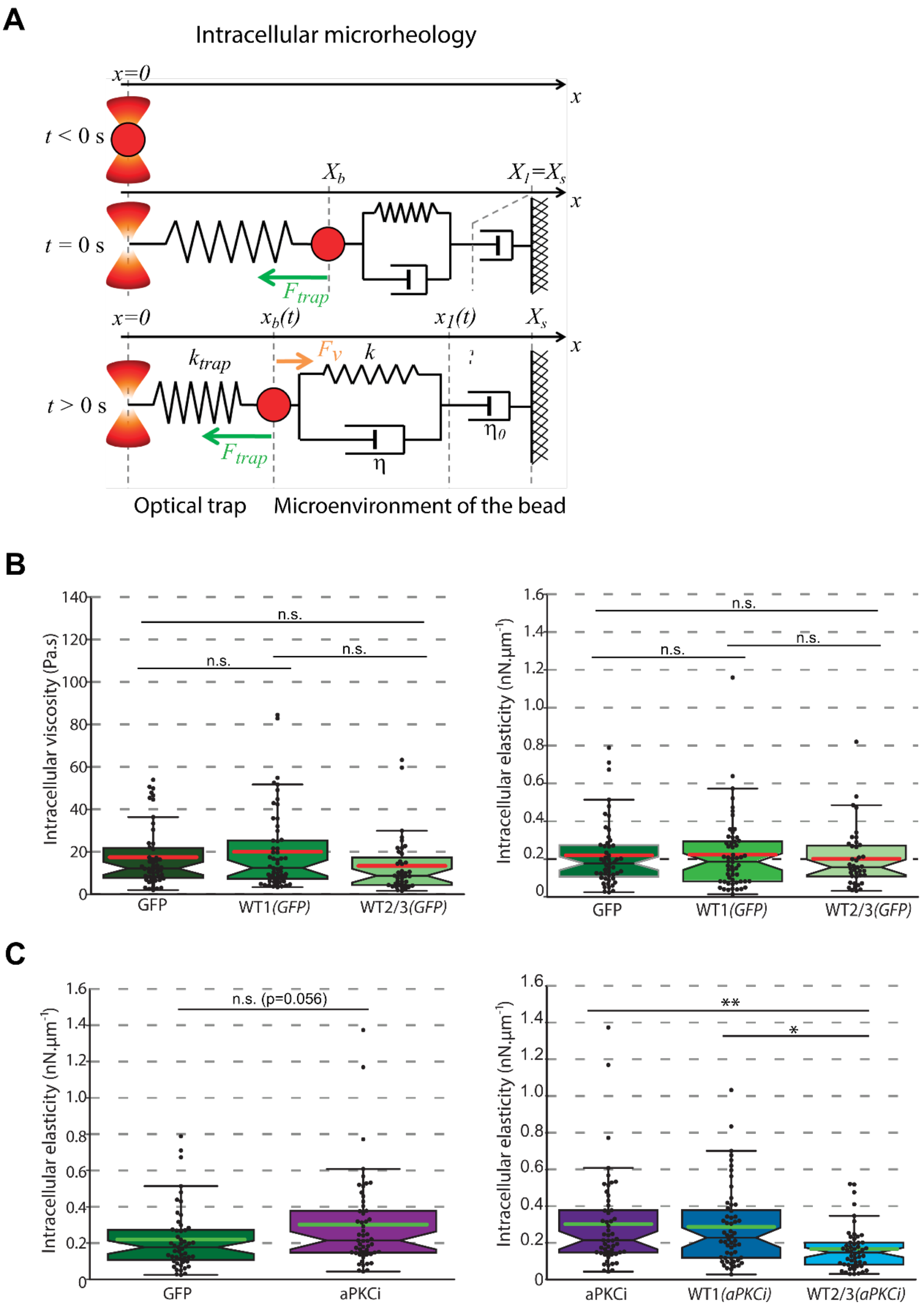

**Fig. S2.**

(related to Fig. 2)

**Optical-tweezers based microrheology to measure intracellular viscosity and elasticity**.

(**A**). Viscoelastic model for intracellular microrheology experiments. The forces acting on an internalized bead trapped in an optical trap at a distance $x_b(t)$ from the trap center are modelled by a spring of stiffness $k_{trap}$ representing the restoring force due to the displacement of the bead from the trap center ($F_{trap} = k_{trap}x_b(t)$) and by a Kelvin-Voigt viscoelastic element (spring of stiffness $k$ and a dashpot of viscosity $\eta$ in parallel) and a dashpot of viscosity $\eta_0$ in series representing the force $F_V$ exerted by the cytoplasm on the bead. At $t = 0$ s, a $X_s = 0.5$ µm step displacement is applied to the stage along the $x$ direction which results in an initial displacement of the bead within the cytoplasm denoted $X_b$. At $t > 0$ s, the bead position relaxes towards the trap center so that $x_b(t) < X_b$.

(**B**). The viscoelastic properties of the cytoplasm are not affected by the presence of GFP cells in a control heterogeneous MCF10-A cell monolayer in which GFP cells are mixed with WT cells. Intracellular viscosity and elasticity measured from viscoelastic relaxation experiments using optical tweezers based microrheology are not statistically different in GFP, WT1(*GFP*) and WT2/3(*GFP*) cells. Data are from n=51, 56 and 39 GFP, WT1(*GFP*) and WT2/3(*GFP*) cells respectively, from three independent experiments. Kruskal-Willis tests were performed; ns, non-significant.

(**C**). The elasticity of the cytoplasm is not affected by overexpression of GFP or aPKCi in heterogeneous cell monolayers. Graphs show the elasticity measured from viscoelastic relaxation experiments in GFP and aPKCi cells (left) and in aPKCi, WT1(*aPKCi*) and WT2/3(*aPKCi*) cells (right). Data are from n=51, 53, 56 and 51 GFP, aPKCi, WT1(*aPKCi*) and WT2/3(*aPKCi*) cells respectively, from three independent experiments. A Student's t-test (left) and a Kruskal-Willis test (right) were performed; *p< 0.05, and **p< 0.01, non-significant if not specified.

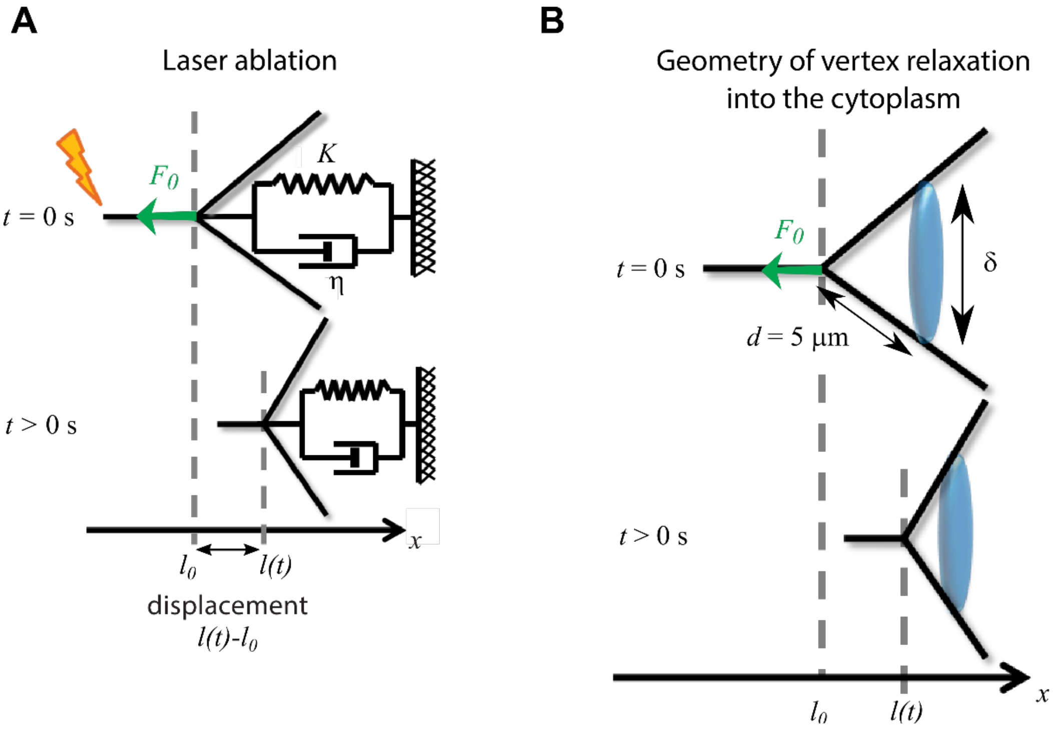


**Fig. S3.**
(related to Figs. 1-4)
**Viscoelastic model used to deduce the junctional tensile force and elasticity at a given vertex from the combination of laser ablation and intracellular microrheology experiments.** A junction is ablated at $t = 0$ s then recoils in the cytoplasm of the neighboring cell ($t > 0$ s).

(**A**). Before ablation, at time $t \leq 0$, the vertex is immobile and the junctional tensile force $F_0$ is balanced by the elasticity $K$ of the other junctions (the vertex elasticity) such that $F_0 = K l_0$, where $l_0$ is the vertex initial position. After ablation, at time $t > 0$, the vertex experiences both the elastic force due to vertex elasticity $K$ and a viscous drag from the cytoplasm modelled as a fluid of viscosity $\eta$, and its position $l(t)$ relaxes.

(**B**). Geometry of the vertex relaxing into the cytoplasm of the neighboring cell. The blue disk of diameter $\delta$ models the vertex and associated cell membranes which relax in the neighboring cell cytoplasm after junction ablation.

### Supplementary Figure 4

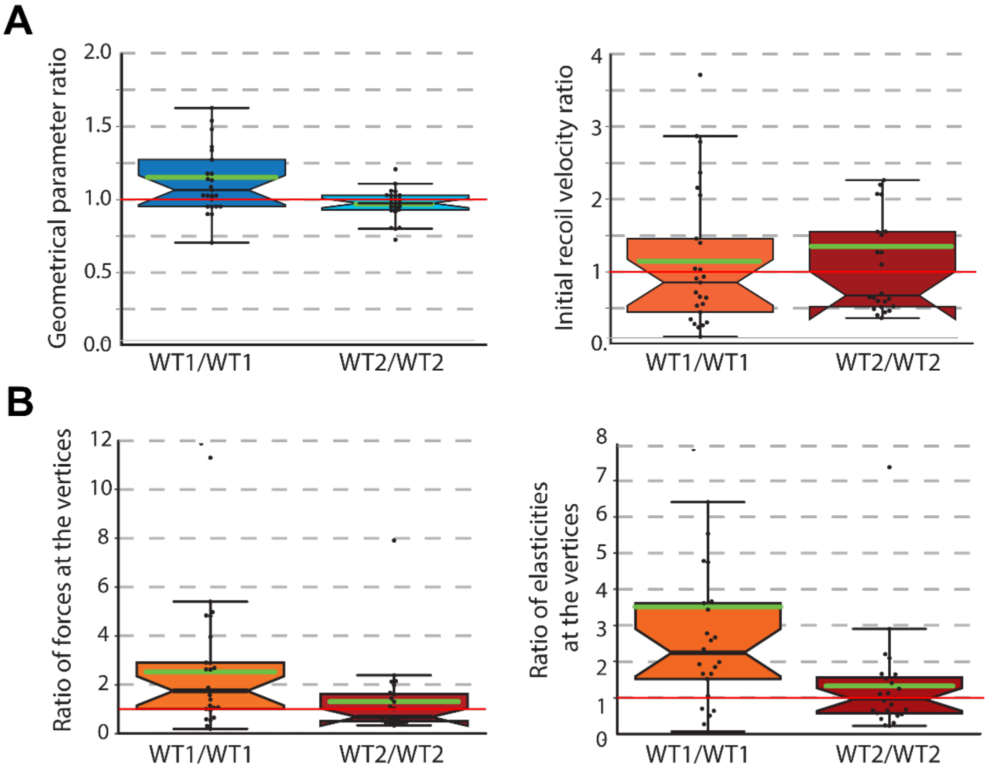


**Fig. S4.**

(related to Fig. 3)

**Characterization of the WT1/WT1 and WT2/WT2 asymmetric junctions.**

(**A**). The two vertices of WT1/WT1 junctions (V1 and V2) and of WT2/WT2 junctions (V4 and V5) have similar geometrical parameters and initial recoil velocities. Graphs show the ratios between the geometrical parameters of the two vertices (left) and the ratios between the initial recoil velocities of the two vertices (right) of WT1/WT1 and WT2/WT2 junctions. Ratios are calculated as the value of the V1 vertex divided by that of the V2 vertex for WT1/WT1 junctions, and as the value of the V4 vertex divided by that of the V5 vertex for WT2/WT2 junctions. Data are from n=25 and 24 WT1/WT1 and WT2/WT2 junctions respectively, from three independent experiments. The red line highlights the symmetric case for which the ratio is equal to one. Note that no statistical test is provided because ratios are compared to one and not between conditions.

(**B**). WT1/WT1 are more asymmetric than WT2/WT2 junctions. Graphs show the ratios between the tensile force at the two vertices (left) and the ratios between the elasticity of the two vertices (right) of WT1/WT1 and WT2/WT2 junctions. Ratios are calculated as the value of the V1 vertex divided by that of the V2 vertex for WT1/WT1 junctions, and as the value of the V4 vertex divided by that of the V5 vertex for WT2/WT2 junctions. Data are from n=25 and 24 WT1/WT1 and WT2/WT2 junctions respectively, from three independent experiments. The red line highlights the symmetric case for which the ratio is equal to one. Note that no statistical test is provided because ratios are compared to one and not between conditions.
